# ManualQC: A MATLAB toolbox for visual EEG quality control and rater calibration

**DOI:** 10.64898/2026.11.20.768044

**Authors:** Yi Xu, Liangkai Sun, Lingzhi He, Zhipeng Cao

## Abstract

ManualQC is a MATLAB toolbox for visual quality control (QC) of epoched electroencephalography (EEG) data. It provides structured QC records of each rater’s selections and comments on epochs, channels, and independent components (ICs). The RaterTool module combines agreement summaries with waveform replay, displaying their epoch and channel markings on the same pre-QC recording for joint review. We illustrate the QC workflow using 20 ERP CORE datasets and demonstrate rater comparison and replay functions using results from three laboratory research assistants on a separate set of EEG recordings. Together, structured QC records and waveform replay facilitate the documentation of manual QC decisions and joint review of disagreements.

**Metadata:** 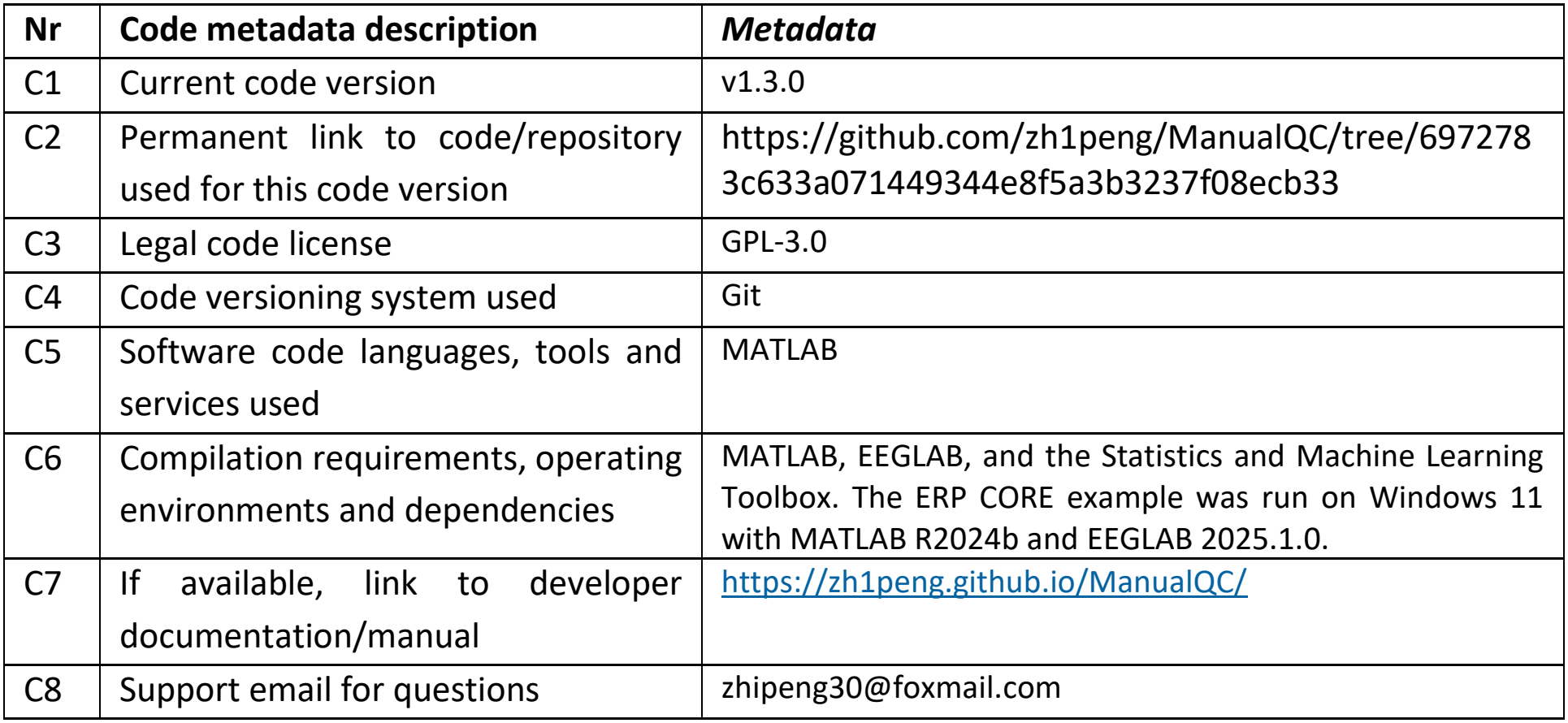

## 1. Motivation and significance

Electroencephalography (EEG) recordings contain artifacts from eye movements, muscle activity, electrode disturbances, and environmental noise. Detecting and reducing these unwanted artifacts is a key step in EEG data processing, and numerous automated methods have been developed for this purpose. For example, FASTER detects statistical outliers in channels, epochs, and independent components (ICs) [1]. Established pipelines such as PREP standardize preprocessing procedures, including robust referencing and noisy-channel detection [2], and ICLabel estimates the probabilities that ICs belonging tobrain or artifact sources [3]. Despite these advances, automated methods may not identify all artifacts reliably. For instance, IC classifications can be uncertain, and components can contain mixed sources [3].

Manual quality control (QC) allows researchers to inspect and remove residual artifacts, but the QC decisions can differ between raters. Raters may select different data for rejection [4], and even expert artifact annotations may differ when the same segments are reassessed [5]. Recording the decisions is therefore important for making EEG preprocessing more transparent and reproducible. However, artifact handling is often insufficiently reported. For example, a review of 132 visual N400 studies published from 1980 to 2018 found that 28.0% described artifact handling only as correction or rejection, while 40.2% did not report the number of trials remained after artifact and/or behavioural exclusions [6]. A simple statement that the data were visually inspected does not reveal which epochs, channels, or ICs were rejected. EEG reporting guidelines and COBIDAS MEEG recommendations also emphasize visual checks of processed data and the reporting of artifact criteria, preprocessing procedures, and retained-data counts [7,8]. Commonly used packages, including EEGLAB [9], FieldTrip [10], and MNE-Python [11] allow users to inspect channel, epoch, and IC properties and manually reject artifacts. However, their processing workflows do not necessarily provide a unified and easy way of recording QC decisions and comments. Researchers may therefore need to manually document and organize the resulting records across datasets.

Another challenge in manual EEG quality control is helping raters consistently apply the laboratory’s artifact criteria. New raters need practice to become familiar with laboratory standards, while experienced raters need opportunities to discuss ambiguous cases. EEG study guidance recommends supervised practice, joint review of recordings, and continuing review to reduce variability across raters and drift in their criteria over time [12]. For training and calibration, raters need to identify disagreements and examine the corresponding signals together. When these tasks are handled separately, researchers may need to compare record files, calculate agreement, and manually locate the relevant signals. A workflow that links the numerical comparisons to visual inspection could make these training and calibration sessions easier to organize. To address these gaps, we developed ManualQC, a MATLAB -based toolbox that supports recording manual QC decisions, structured documentation of QC decisions, and comparison of QC decisions across raters.

## 2. Software description

ManualQC is developed in MATLAB and uses EEGLAB EEG data structure and functions for visualization, processing, and saving. It can be launched as an EEGLAB plugin or run directly in MATLAB after EEGLAB paths have been configured.

### 2.1. Software architecture

ManualQC consists of two main components (Fig. 1): a QC workbench and RaterTool module. The workbench allows raters to inspect and process one EEG dataset at a time and saves the rater’s decisions in the processed dataset and in a CSV summary file. RaterTool matches QC records from different raters by dataset and displays their selections on the same pre-QC recording.

**Fig. 1.**
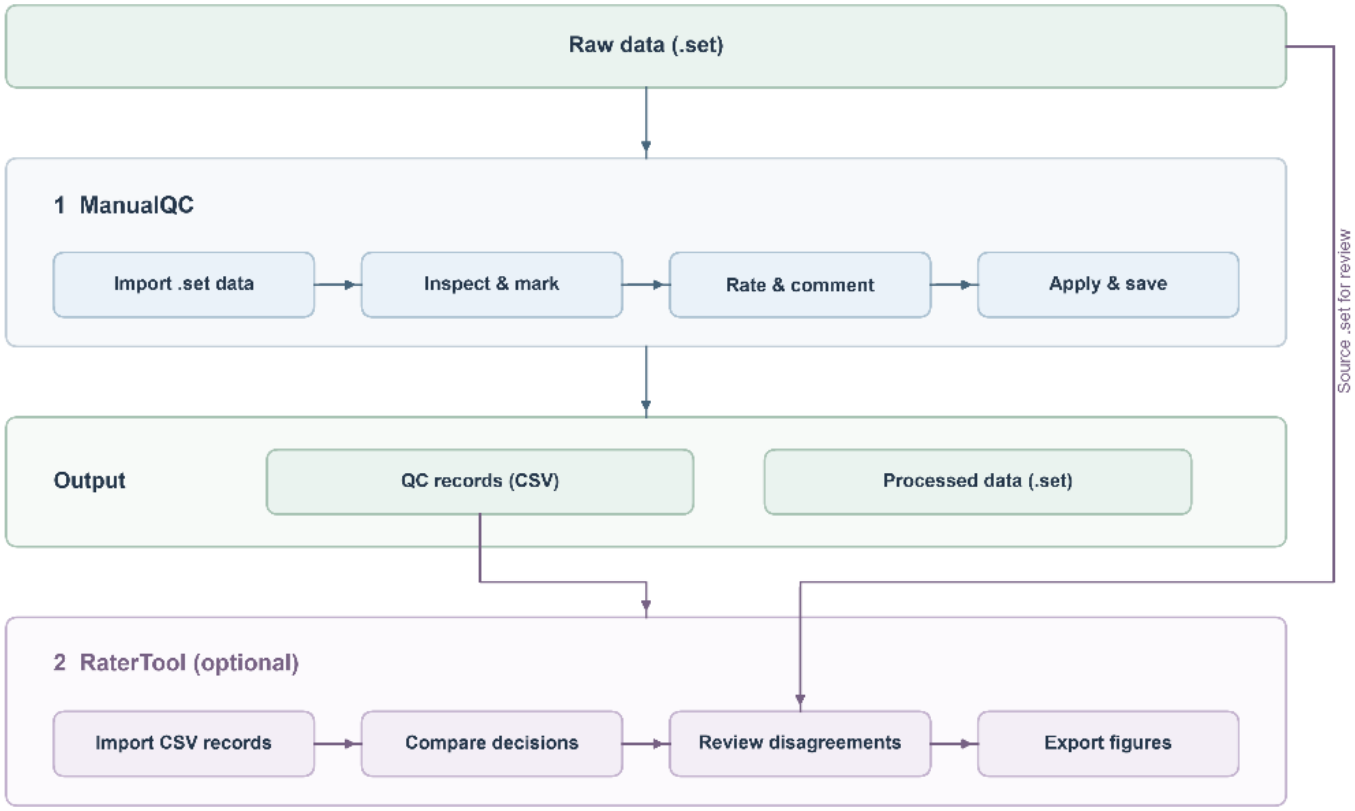
Overview of the ManualQC workflow. Raters inspect and process datasets in the QC workbench, which saves their decisions in each dataset and in CSV summaries files. RaterTool uses these records to compare raters’ selections and display their epoch and channel markings on the same recording for group review and training.

### 2.2. Software functionalities

#### 2.2.1 Dataset selection and inspection

ManualQC searches a user-specified directory for EEGLAB ‘.set’ files using a regular expression. After loading a dataset, ManualQC displays its basic information. Users can handle the detected datasets sequentially or enter an index to start with a specific dataset.

ManualQC supports manual inspection of epochs, channels, and ICs. For epochs inspection, users open the EEGLAB waveform viewer (eegplot) [9] to mark bad epochs for exclusion. For channels inspection, users review channel locations, single-trial voltage plots, average waveforms across epochs, and power spectra to mark bad channels. For ICs inspection, users inspect component scalp topographies activity time courses, and power spectra before selecting components for removal. Decisions about artifact status are made by the user, and the software provides the tools for visual inspection, annotation, and processing workflow.

These selections are applied when saving the dataset. ManualQC first removes the selected ICs, then rejects the marked epochs and finally applies spherical interpolation to the marked bad channels. ManualQC then saves the processed dataset and exports the QC records to a CSV summary file. Alternatively, raters can inspect the signals after IC removal before deciding which epochs to reject or which channels to interpolate. The optional “Remove Bad ICs First” command removes the selected ICs to the recording in memory. Once confirmed, the command clears all existing epoch and channel markings. Raters need to reassess the updated waveforms before marking epochs or channels again.

#### 2.2.2 Dataset assessment

ManualQC provides two forms of dataset-level quality information. The manual rating reflects the rater’s overall assessment of dataset usability, whereas the ‘Data Quality value’ measures amplitude variability before and after QC. ManualQC automatically calculates the ‘Data Quality value’ as a summary measure of signal-amplitude variability:

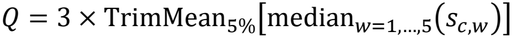

To calculate this value, ManualQC selects up to 5,000 points at approximately equal intervals along the time dimension and divides them into five equal-length windows. For each window, the toolbox calculates the standard deviation of each channel. The median of the five window-specific standard deviations is calculated for each channel, ManualQC then calculates the 5% trimmed mean across channels, and multiplies the resulting value by three. The across-window median and cross-channel trimmed mean may reduce the influence of extreme values. However, this scalar may be insensitive to brief artifacts and can also yield low values for a flat signal. Rounded scores of ≤18, 19–28, and ≥29 are categorized as high, moderate, and low quality, respectively. The Data Quality value only serves as a supplementary measure, while the final assessment of dataset usability is based on the rater’s inspecting the recording. After inspecting the dataset, the rater assigns one of three ratings: ‘Usable’, ‘Recheck’, or ‘Not Usable’. The corresponding filename prefixes are ‘qced_’, ‘recheck_’, and ‘bad_’, respectively. The output filenames retain the original participant and task identifiers. Raters can explain the reasons for their decisions in the comments field.

In addition to the manual rating,

#### 2.2.3 QC records and cross-dataset management

ManualQC stores QC information in the ‘EEG.etc.manualqc’ structure of each processed dataset. The stored informationinclude the dataset identifier (dataset_name), rejected-epoch records, interpolated channel names, removed IC indices, rating, comments, and ‘Data Quality value’ before and after QC. Storing these fields with the processed dataset allows users to review the recorded QC decisions alongside the data.

When epochs are rejected, ManualQC stores both the indices and their indexing convention. The ‘rejected_epochs_current’ field stores epoch indices in the dataset immediately before rejection. The ‘rejected_trials_id’ field stores the original trial identifiers when a mapping is available,or the current epoch indices otherwise. The ‘rejected_id_space’ field indicates which indexing convention is used. In addition to records within individual datasets, ManualQC maintains a cross-dataset summary in the MATLAB workspace variable ‘QC_log’. After each dataset is saved, the accumulated records are written to a CSV backup file. A final summary is generated when the last dataset in the processing list is saved. Records from different QCsessions can also be combined for each rater for subsequent comparison, provided that dataset identity and indexing conventions remain consistent.

#### 2.2.4 RaterTool: comparing records across raters

RaterTool is a separate tool that compares QC records created by different raters for the same set of datasets. It imports one CSV or XLSX file per rater, matches records by dataset identifier, and compares epoch rejection, channel interpolation, and IC removal. For each dataset, overlap is calculated across the raters who recorded decisions for it. Datasets with only one rater remain in the summary with no agreement estimate.

Agreement in QC markings is assessed by different raters overlap. For two raters, the Jaccard index is the number of items both marked divided by the number marked by either of them [13]. RaterTool reports the average over all rater pairs for each dataset. Group Jaccard divides the number of items marked by every rater by the number marked by at least one rater. Precision is the proportion of the first rater’s markings that the second rater also made. Recall is the proportion of the second rater’s markings that the first rater also made. The order follows the pairwise table. F1 is their harmonic mean and, like Jaccard, is independent of rater order.

When EEG metadata are enabled, the shared pre-QC dataset defines the available epochs, channels, and ICs. Each item is coded as marked or unmarked, and Cohen’s κ (two raters) or Fleiss’ κ (three or more) is computed [14,15]. A κ estimate is not reported if a marking cannot be mapped to an item or if only one category for valid κ calculation is observed. QC_Before and QC_After are treated separately. For each continuous score, RaterTool computes the two-way random-effects, absolute-agreement ICC(2,1) across datasets with a finite score from every rater and reports the number of contributing datasets [16].

These comparisons require consistent dataset identifiers, dataset versions, and item-index conventions. By convention, two empty marking sets have Jaccard and F1 equal to 1, so the overlap values should be read together with the number of markings actually recorded. The CSV summary stores current epoch positions. When the pre-QC EEG contains an ‘EEGdojo.TrialID’ mapping, the epoch κ calculation interprets the imported values as original trial identifiers. Users should check that the exported positions and original identifiers coincide for the marked epochs.

For replay, RaterTool uses the current epoch positions to display the recorded selections on the shared pre-QC waveforms. Colored bands show the epochs each rater marked for rejection, and colored triangles beside the channel labels identify channels selected for interpolation. Raters can inspect the signal waveforms and open channel or IC properties to review a disagreement.

## 3. Illustrative examples

### 3.1 Preparation of the example datasets

The demonstration used 20 participants (sub-001 to sub-020) from the ERP CORE P3 dataset [17]. Before manual QC, each dataset contained 30 scalp EEG channels, 200 epochs, and 29 ICs, yielding 4,000 epochs across all datasets. Testing was performed on Windows 11 with MATLAB R2024b, EEGLAB 2025.1.0, and ManualQC v1.2.0. Preprocessing included resampling, 1–40 Hz filtering, and average referencing. Extended Infomax ICA was then performed on the continuous data, with PCA dimensionality set according to the rank of the average-referenced data. Epochs were extracted from -0.2 s to 0.8 s relative to stimulus onset. No epoch-wise baseline correction was applied. Before manual QC, all 200 epochs, 30 selected scalp channels, and 29 ICs were retained. No artifact-contaminated epochs or components had been removed.

### 3.2 Inspecting and saving an example dataset

The filename expression ^Final_.*\.set$ selects the prepared datasets in the workbench (Fig. 2A). In sub-001, the rater marked epochs 93 and 94 for rejection after observing conspicuous abnormal oscillations (Fig. 2B). The rater selected no channels for interpolation (Fig. 2C), marked ICs 1, 4, 5, and 8 for removal (Fig. 2D), and rated the dataset ‘Usable’.

**Fig. 2.**
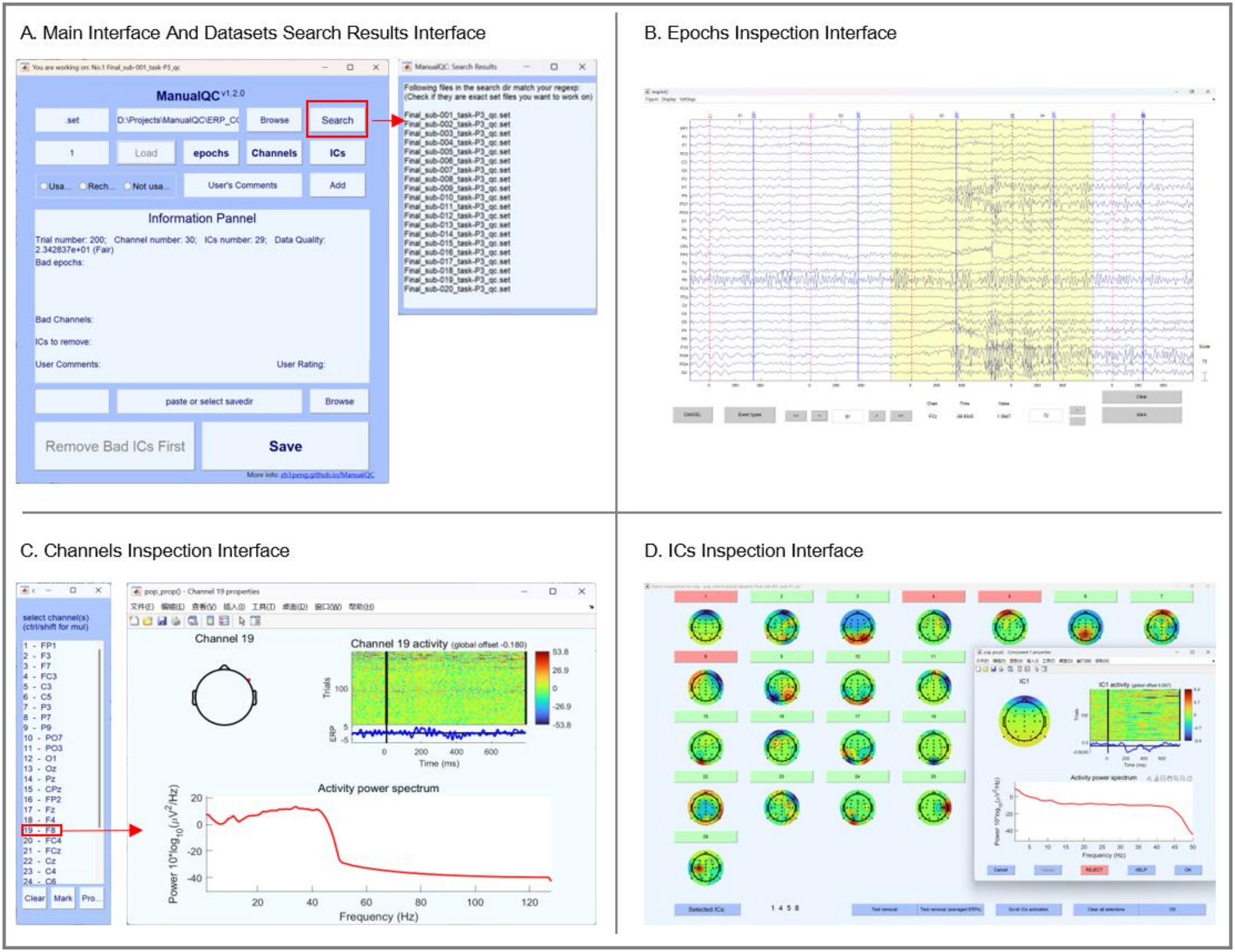
Visual inspection of sub-001 in ManualQC v1.2.0. (A) Workbench and search results. (B) Waveforms with the marked epochs. (C) Channel list and properties. (D) IC topographies and component properties.

### 3.3 QC outcomes and record consistency

Across the 20 datasets, the rater removed 44 ICs and rejected 11 epochs from five datasets, leaving 3,989 epochs (Table 1). All datasets retained 30 channels and were rated ‘Usable’. No channels were selected for interpolation, and all comments fields contained ‘None’.

**Table 1.**
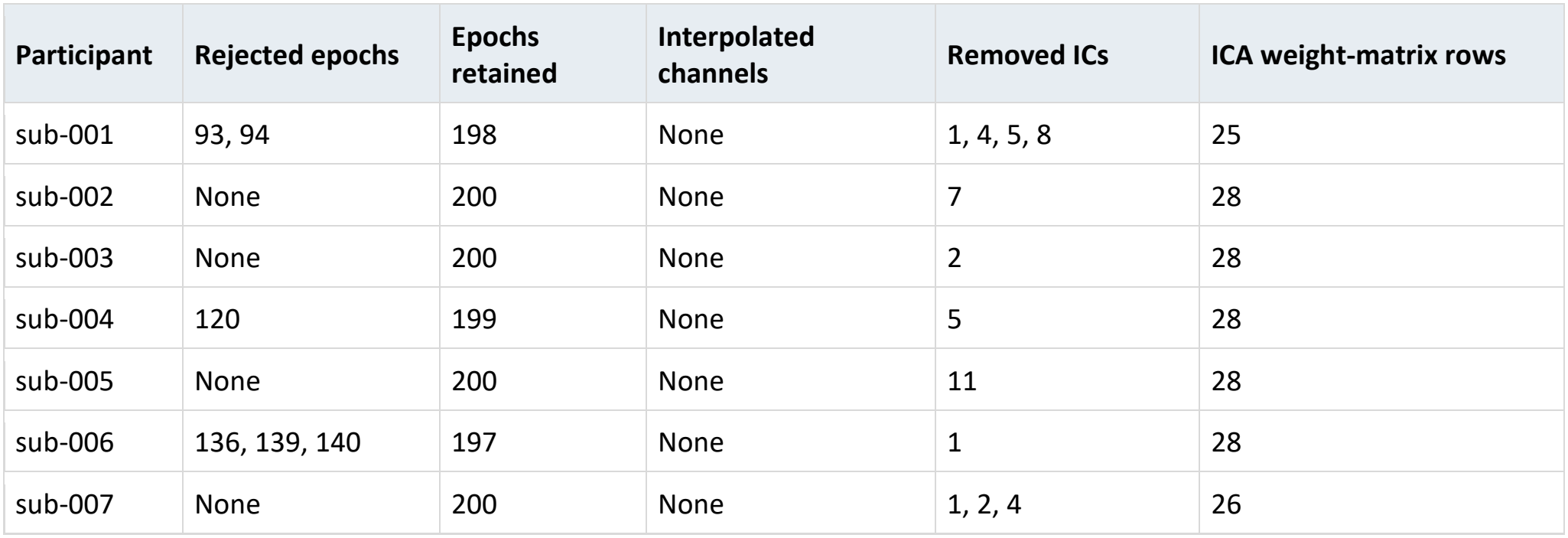
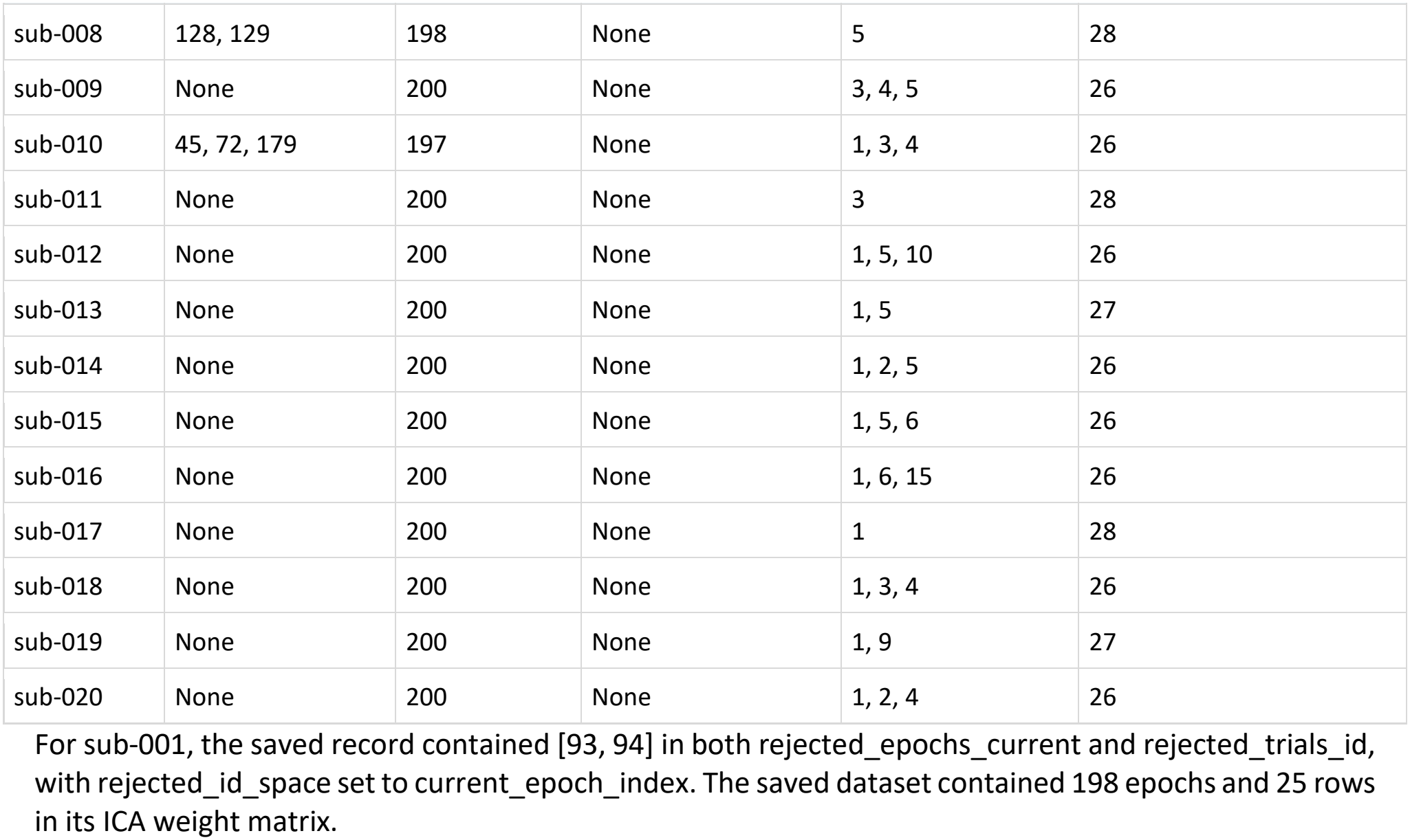
Manual QC selections and saved data dimensions in the ERP CORE example.

| **Participant** | **Rejected epochs** | **Epochs retained** | **Interpolated channels** | **Removed ICs** | **ICA weight-matrix rows** |
|---|---|---|---|---|---|
| sub-001 | 93, 94 | 198 | None | 1, 4, 5, 8 | 25 |
| sub-002 | None | 200 | None | 7 | 28 |
| sub-003 | None | 200 | None | 2 | 28 |
| sub-004 | 120 | 199 | None | 5 | 28 |
| sub-005 | None | 200 | None | 11 | 28 |
| sub-006 | 136, 139, 140 | 197 | None | 1 | 28 |
| sub-007 | None | 200 | None | 1, 2, 4 | 26 |
| sub-008 | 128, 129 | 198 | None | 5 | 28 |
| sub-009 | None | 200 | None | 3, 4, 5 | 26 |
| sub-010 | 45, 72, 179 | 197 | None | 1, 3, 4 | 26 |
| sub-011 | None | 200 | None | 3 | 28 |
| sub-012 | None | 200 | None | 1, 5, 10 | 26 |
| sub-013 | None | 200 | None | 1, 5 | 27 |
| sub-014 | None | 200 | None | 1, 2, 5 | 26 |
| sub-015 | None | 200 | None | 1, 5, 6 | 26 |
| sub-016 | None | 200 | None | 1, 6, 15 | 26 |
| sub-017 | None | 200 | None | 1 | 28 |
| sub-018 | None | 200 | None | 1, 3, 4 | 26 |
| sub-019 | None | 200 | None | 1, 9 | 27 |
| sub-020 | None | 200 | None | 1, 2, 4 | 26 |

For sub-001, the saved record contained [93, 94] in both rejected\_epochs\_current and rejected\_trials\_id, with rejected\_id\_space set to current\_epoch\_index. The saved dataset contained 198 epochs and 25 rows in its ICA weight matrix.

We checked the input and output files separately in EEGLAB to verify the functionalities of ManualQC workbench for recorded QC selections. For each dataset, the decrease in the number of epochs and in the number of ICA weight-matrix rows matched the epoch rejections and IC removals recorded in the CSV summary.

### 3.4 Rater comparison and replay

Three research assistants assessed a separate collection of EEG recordings acquired with 64-channel Neuracle or 128-channel EGI systems. We imported their logs (rater_czr.xlsx, rater_hlz.xlsx, and rater_pcj.csv) into RaterTool. Ten recordings had QC records from all three assistants (Fig. 3A). Across these ten recordings, mean pairwise Jaccard was 0.38 for rejected epochs, 0.46 for interpolated channels, and 0.40 for removed ICs. The corresponding mean group Jaccard values were 0.23, 0.32, and 0.33. In three recordings, no rater marked an IC, resulting in undefined IC κ values. Excluding them reduced mean pairwise IC Jaccard to 0.14. Within each recording, all three raters had identical logged pre-QC scores. The post-QC scores stored in the logs had an ICC(2,1) of 0.93 across the ten recordings. Channel κ was undefined in the two recordings for which no rater marked a channel. We replayed MID task EEG of sub-165. The bundled pre-QC file contains 59 channels and 327 epochs. With all three raters selected, we set the starting epoch to 175 (Fig. 3B). All three retained epoch 175 and rejected epochs 177 and 178. Only rater_czr and rater_pcj rejected epoch 176. Only rater_pcj rejected epoch 179.

**Fig. 3.**
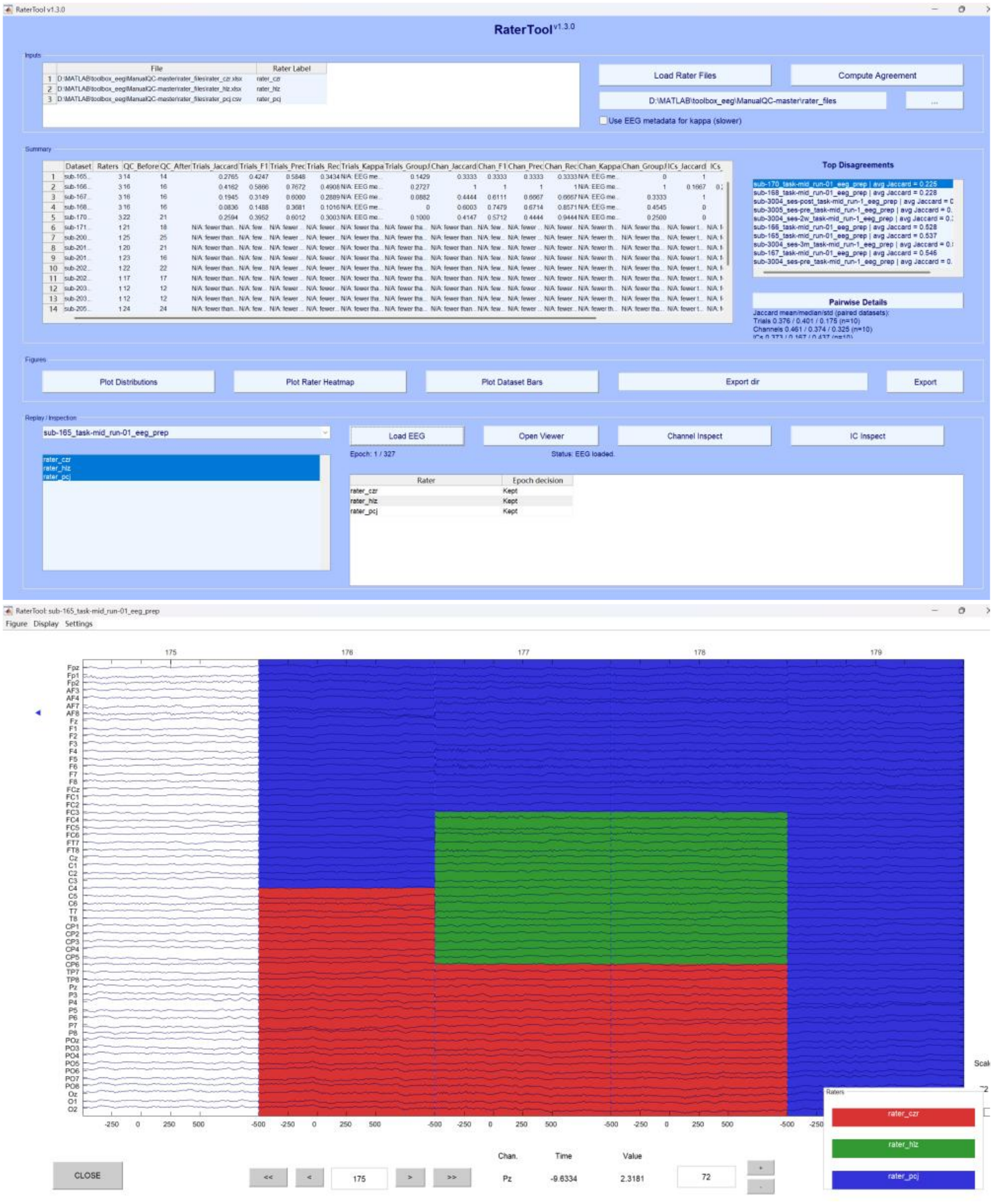
Rater comparison and replay in RaterTool v1.3.0. (A) Agreement summary and replay controls with sub-165 loaded. The QC score columns show values from the original rater logs. (B) Pre-QC waveforms for epochs 175–179. The counts above the viewer give the number of raters who rejected each epoch. Red, green and blue bands identify rater_czr, rater_hlz and rater_pcj, respectively. The blue triangle marks AF8, selected by rater_pcj for interpolation.

## 4. Impact

ManualQC supports structured documentation of manual QC by recording each rater’s selections and comments for individual EEG datasets. These records reduce the need for separate documentation and provide a basis for reviewing QC decisions and reporting artifact handling procedures. They can complement reporting templates such as ARTEM-IS [22] by providing dataset-level information about the selections. ManualQC was developed from EEGQC_GUI (https://github.com/zh1peng/EEGQC_GUI), an earlier laboratory interface developed for manual QC after automated preprocessing. ManualQC has been used in our previous studies supporting visual inspection of EEG data [18-21]. For example, it was used to inspect EEG data in a study of reward feedback processing in patients receiving methadone maintenance treatment [21]. These studies demonstrate its practical use in EEG research, although its adoption has so far been largely limited to laboratories involved in its development. Through this publication, we aim to introduce ManualQC to the broader EEG research community and encourage its wider use.

The RaterTool module of Manual QC also supports rater training and calibration by combining agreement summaries with waveform replay, allowing raters to identify disagreements and jointly review the corresponding EEG signals. RaterTool display each rater’s epoch and channel markings on the shared pre-QC recording. Rather than comparing QC record files and then manually locating the corresponding data, researchers can inspect the records alongside the data. This replay functionality provides a practical tool for the joint review recommended in EEG study guidance [12], giving trainees and experienced raters a shared basis for discussing difficult cases. We have used ManualQC to train multiple raters within our laboratory. We hope that it will also benefit other laboratories by supporting efficient, structured training in manual EEG QC.

## 5. Conclusions

ManualQC integrates visual quality control of epoched EEG data with structured QC records of each rater’s selections and comments. Its RaterTool module provides agreement summaries with waveform replay, allowing researchers to review different raters’ epoch and channel markings on the same pre-QC recording. Together, these features support the documentation of manual QC decisions and joint review of disagreements, providing a practical basis for discussing and clarifying artifact criteria.

## Data and software availability

The ERP CORE P3 data used in the processing example are available under CC BY-SA 4.0 at https://doi.org/10.18115/D5JW4R [17]. The ManualQC source code is available under GPL-3.0 at https://github.com/zh1peng/ManualQC/.

## CRediT authorship contribution statement

Yi Xu: Writing – original draft, Visualization. Liangkai Sun: Writing – review & editing. Lingzhi He: Writing – review & editing. Zhipeng Cao: Conceptualization, Methodology, Formal analysis, Writing – review & editing, Visualization, Supervision, Funding acquisition.

## Declaration of Generative AI and AI-assisted technologies in the writing process

During manuscript preparation, the authors used a large language model to refine and proofread the text and broaden the literature search. The authors reviewed and edited all AI-generated content as needed and take full responsibility for the content of this publication.

